# Imaging analysis to quantitate the Interplay of membrane and cytoplasm protein dynamics

**DOI:** 10.1101/2021.12.01.470244

**Authors:** N Kislev, M Egozi, D Benayahu

## Abstract

Plasma membrane proteins are extremely important in cell signaling and cellular functions. Protein expression and localization alter in response to various signals in a way that is dependent on cell type and niche. Compartmental quantification of the expression of particular proteins is a very useful means of understanding their role in cellular processes. Immunofluorescence staining is frequently used to investigate the distribution of proteins of interest. Here, we present an imaging method for quantifying the membrane to cytoplasm ratio (MCR) of proteins analyzed at single-cell resolution. This technique provides a robust quantification of membrane proteins and contributes new insights into membrane expression dynamics. We have developed a protocol that uses immunostaining to assess protein expression according to the fluorescent cellular distribution and to compute the MCR. The method was applied to measure the MCR of glucose transporter 4 (GLUT4) in response to insulin in 3T3-L1 cells, an in-vitro model for adipocyte function and adipogenesis. The results revealed informative changes in the subcellular localization of GLUT4 following insulin induction. MCR analysis is a powerful imaging tool that can be generally applied to membrane proteins to provide a rapid and efficient quantitative analysis of protein distribution and sub-cellular processes in cells.

## Introduction

The plasma membrane (PM) is composed of a semipermeable lipid bilayer consisting of phospholipids that separates the cell from the surrounding environment. The structure of the PM is fundamental to various cellular processes and provides separation and physical support to the cell while the membrane proteins administrate the transportation of nutrients and interaction with other cells and the environment. Each cell type has a unique composition of PM proteins depending on the specific niche, interactions, and cellular/ metabolic state. Both integral membrane proteins that penetrate the PM, and peripheral proteins, are vital components that allow the PM to communicate with the niche via receptors that attach the cells, and others that mediate the cellular response to various factors and activate signaling pathways. Differential expression of membrane proteins is dynamic and has implications for cell function and specifically on processes such as proliferation, migration, cell death, and differentiation. The importance of detecting and quantifying alterations in expression levels and distribution of PM proteins is at the foreground of cell and molecular biological research^1–4^.

Adipogenesis is the differentiation process of pre-adipocytes/fibroblasts (FB) to mature adipocytes (AD). This is a dynamic and complex process that involves the regulation of a variety of differentiation factors and co-factors^5^, and is accompanied by morphological changes as the spindle-like FBs become rounder and accumulate lipid droplets. Differentiated AD exhibit a membrane protein profile that differs from that of FB cells, reflecting their specialized function^6,7^. For example, the insulin dependent glucose transporter, GLUT4, is a key protein that is upregulated in adipocytes. Insulin is crucial for adipocyte function and adipogenesis. The hormone induces a significant signaling pathway in adipocytes^8–10^ by binding to the insulin receptor (IR) and activating a pathway involving the regulation and translocation of GLUT4-containing cytoplasmic storage vesicles to the PM. This enables the uptake of glucose into the cell, which is required to maintain metabolic homeostasis and control the systemic glucose level. A reduction in GLUT4 translocation and docking to the PM is associated with insulin resistance and diabetes^11–13^. Monitoring the shuttling of GLUT4 is vital to understanding the metabolic status of adipocytes and their response to various stimuli^14,15^.

PM protein localization and distribution can be studied using a variety of imaging techniques, although the results are primarily qualitative. In contrast, the method presented here enables a robust and rapid quantification of the PM protein intensity at the single-cell level. ImageJ is used to fractionate images of immunofluorescent (IF) stained cells into their various cellular compartments and to calculate the fluorescence intensities and ratios for each cell. The result allows us to quantify the alterations in the intensity of membrane proteins induced by a multitude of different stimuli and conditions. In the current study, we describe a protocol for quantifying the membrane to cytoplasmatic ratio (MCR) of protein expression in single cells that can analyze individual cell responses even in a heterogeneous population. Here, we applied it to the 3T3-L1 cell line, which is an in-vitro model for adipocyte studies and were able to demonstrate differences in the expression of GLUT4 in AD and FB present in the same culture. Cultured cells were analyzed under induced/basal conditions to monitor the response of the AD to the addition of insulin. Ultimately, the imaging analysis presented here will allow us to quantify the dynamic interplay of GLUT4 expression in the membrane and cytoplasm. Importantly, this method for detection and measurement of the variation in membrane proteins can be applied generally to a variety of cell types. This powerful tool simultaneously investigates molecular regulation networks and single-cell characteristics and can be of great use in advancing our understanding of diverse cellular mechanisms.

## Methods and materials

### Cell Cultures

Mouse embryonic 3T3-L1 pre-adipocytes (American Type Culture Collection) were used as previously described ^16,17^. For the insulin induction, differentiated 3T3-L1 cells were incubated in a Dulbecco’s modified Eagle’s medium without glucose for one hour (Biological Industries, Kibbutz Beit-Haemek, Israel), and then replaced with a growth medium with and without 5 ug/ml insulin for 30 minutes.

### Immunofluorescence staining

Cells were cultured on coverslips, and at the end of the experiments they were fixed in 4% paraformaldehyde containing 30mM sucrose in PBS for 10 minutes at room temperature (RT). For the staining, cells were permeabilized with 0.5% Triton in 1% TBST for 10 minutes and then blocked with a blocking solution (1% TBST containing 1–2% normal goat serum and 1% BSA) and stained with anti-GLUT4 (SC-53566), and then with secondary antibody, Cy3-anti-mouse (Jackson). The stained coverslips were mounted on slides with Fluoroshield™ mounting medium containing 4’, 6-diamidino-2-phenylindole (DAPI). Images were acquired by a Leica SP8 confocal microscope (Leica, Wetzlar, Germany).

### Image processing and data analysis

All steps of the image analysis were performed using FIJI ImageJ software

### Membranal/cytoplasmatic ratio (MCR) measurements

#### Creating the rim boundaries

Figure 1 presents the analysis flow; the outer membrane, inner membrane, and perinuclear region boundaries were marked to analyze the compartmental expression. First, the outer cellular membrane was marked, and the surroundings of the selected cell were cleared (1). Then, the image prepared for thresholding by creating Gaussian Blur until the shape was uniform (2). This was followed by thresholding the image at the point in which the cell has a complete membrane (3). The distinguished cell membrane was marked as the extracellular side of the membrane. (4). Afterwards, the image was eroded to delineate the intracellular side of the cell membrane. (5). Each ring was saved to the region of interest (R.O.I.), forming a membrane rim. The marking accuracy was assessed with the original images (Fig. 1).

**Fig. 1.**
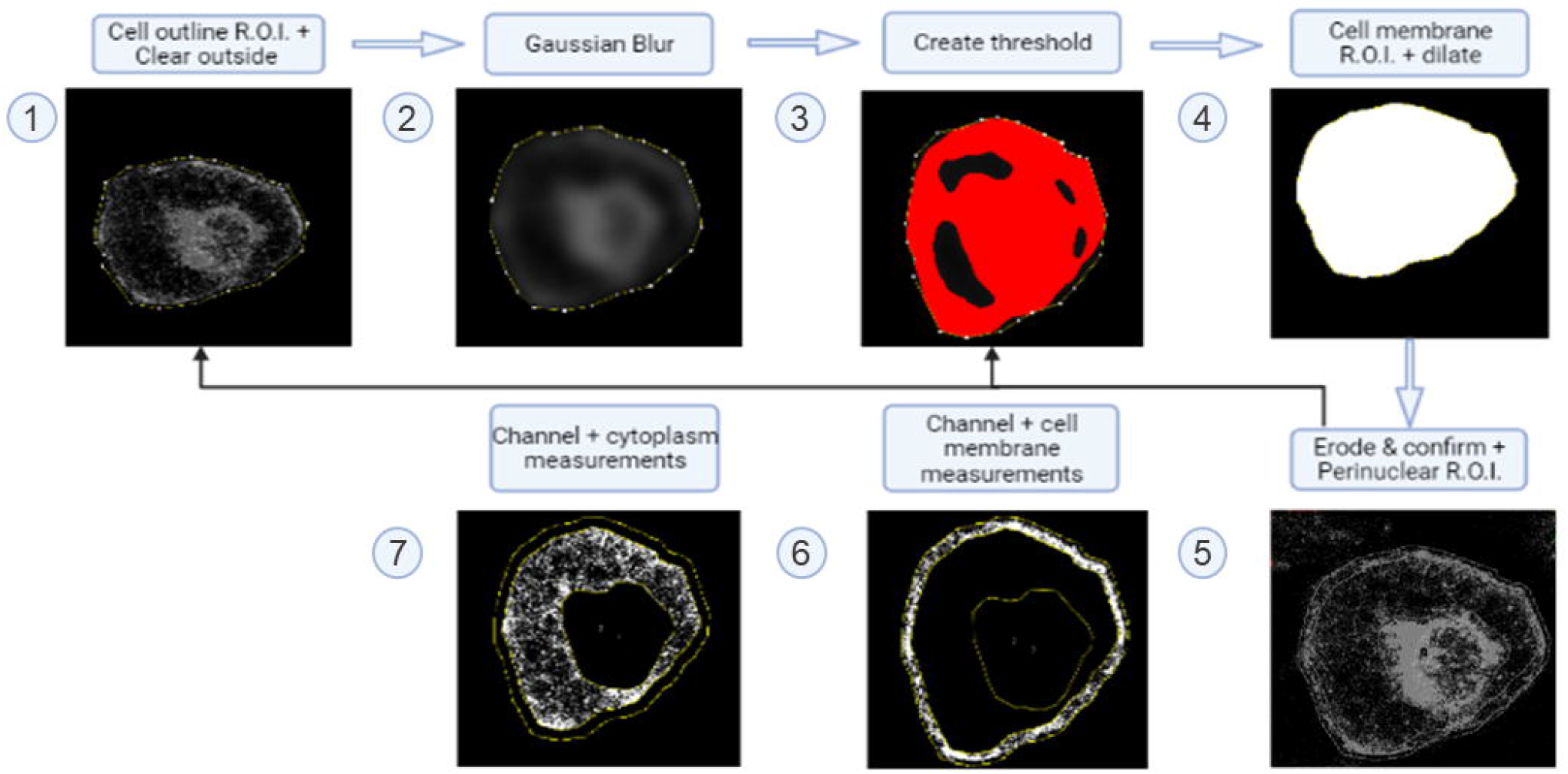
Membranal/cytoplasmatic ratio (MCR) workflow. (1) Selection of the region of interest and clearance of the surroundings (2) Gaussian blur as a prerequisite step for a filled-up shape (3) Thresholding and clearance of the internal shape (4) Dilation of the shape to obtain the extracellular border (5) Erosion of the shape to obtain the intracellular border and a rim is obtained, confirm the accuracy of the membrane rim (if needed repeat the steps above) (6) Clearance of the intracellular filling and membrane intensity measurement (7) Clearance of the nuclear area and cytoplasm intensity measurements.

#### Measurement

The original images were reselected. Then, the outer membrane R.O.I. and the inner membrane R.O.I. were applied to the original image and the outside (background) and the inside (intracellular) were cleared. Next, only the cell membrane was selected in the field of view (FOV) and the intensity of the membrane rim was measured (6). The original image for measurement was uploaded once again. Then the inner membrane R.O.I. was applied to the original image, and the outside and nuclear areas were cleared, and the measurement of the cytoplasm was taken (7) (Fig. 1).

#### Intensity parameters

i. Membranal mean intensity: Measures the mean intensity of the membranal ring. Contains the raw integrated density of the membranal rim fraction divided by the rim area. (The outer membrane area minus the inner membrane area)
ii. Cytoplasmatic mean intensity: Measures the mean intensity of the cytoplasm. Contains the raw integrated density of the cytoplasm fraction divided by the cytoplasmic area.
iii. Membranal/cytoplasmatic ratio (MCR): The ratio of the membranal intensity and the cytoplasmatic mean intensity.

The measurements were taken for visible adipocytes in different fields and cultures presented as mean and standard errors (SEM) of each single parameter. The single-cell analyses were performed to examine the scattering of each parameter.

### Statistical analysis

Statistical analyses were performed by GraphPad Prism v.8.1.1. Results are presented as means ± SEM. All results were tested for normal distribution by Kolmogorov-Smirnov test, and outliers were identified using the ROUT method. Statistical differences comparing the mean values were tested using two-tailed, unpaired, t-tests. A value of p < 0.05 was considered statistically significant.

## Results

### Analysis of cell differentiation and GLUT4 expression at single cell resolution

The MCR method was developed to measure GLUT4 membrane expression in 3T3-L1 cells. As was previously shown by our group^16,17^, there are significant morphological differences during adipogenesis and differentiation of 3T3-L1 cells. Undifferentiated cells (FB) have a spindle-like shape with cytoplasmic processes while differentiated (AD) cells are round and filled with lipid droplets. As differentiation progresses, the accumulating lipid droplets eventually fuse, generating larger and larger lipid droplets in the adipocyte. The cells also differ in their metabolic and molecular functions, as AD specialize in accumulating lipids and taking up glucose through transporters like GLUT4 (Fig. 2A). We examined the expression of GLUT4 in both cell types (FB and AD) in the same culture during differentiation in the presence of basal insulin. The immunofluorescence staining revealed that adipocytes have high levels of GLUT4 storage vesicles, particularly in the perinuclear area. In contrast, the GLUT4 fluorescence in FB is weak and evenly distributed throughout the cell. Figure 2B presents the results, both as stained cells and as the corresponding plots of the immunostaining measurements.

**Fig. 2.**
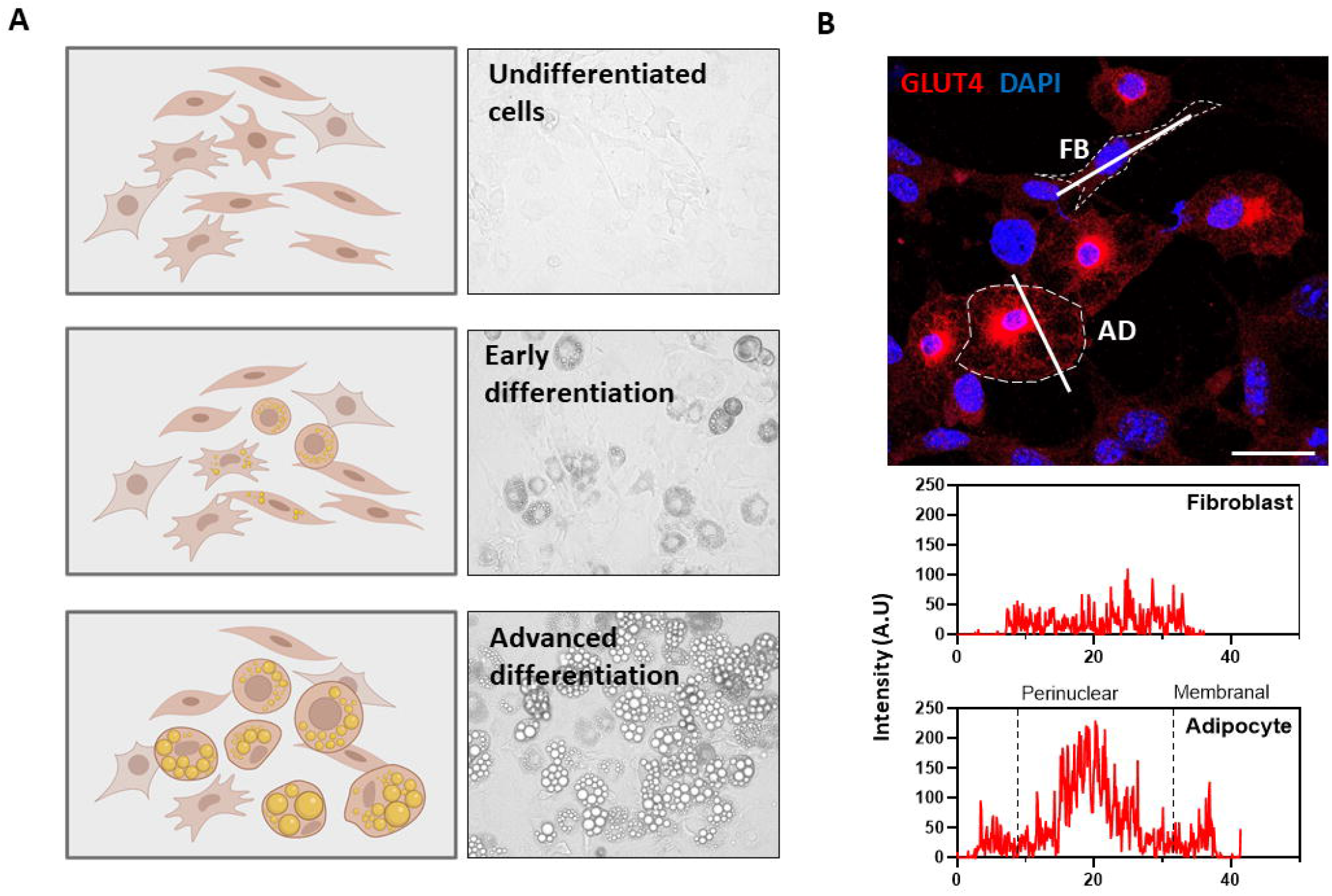
GLUT4 expression in 3T3-L1 cells. (A) Schematic illustration and phase-contrast images along differentiation phases of adipogenesis (Magnification of X200, scale bar=50μm) (B) Immunofluorescence staining of GLUT4 (red), and Nuclei (blue) in differentiated 3T3-L1 cells. (Magnification of X630, scale bar=50μm), Intensity line of GLUT4 level at the membrane and cytoplasmic sites.

### Insulin induction alters the cellular distribution of GLUT4 as measured by the membrane to cytoplasm ratio (MCR)

As shown in Fig. 3, GLUT4 is expressed in AD in cytoplasmic storage vesicles. When stimulated with insulin, the immunostaining indicates that the GLUT4 containing vesicles translocate to the PM (Fig. 4A). The intensity plots reveal a similar pattern, with GLUT4 uniformly distributed in the unstimulated cells but with a distinct membrane and perinuclear expression in insulin stimulated cells (Fig. 4B) and limited intensity in the cytoplasm. MCR calculations according to the method described in Figure 1 indicate a substantially higher membrane intensity in the insulin induced AD than the mostly uniform distribution in FB cells. This translates to an upregulation of 30% in the PM localization of insulin induced cells compared to the non-induced cells (Figs. 4C, 4D). These results demonstrate the potential of single-cell analysis in quantifying the expression and behavior of different membrane proteins under various conditions.

**Fig. 3.**
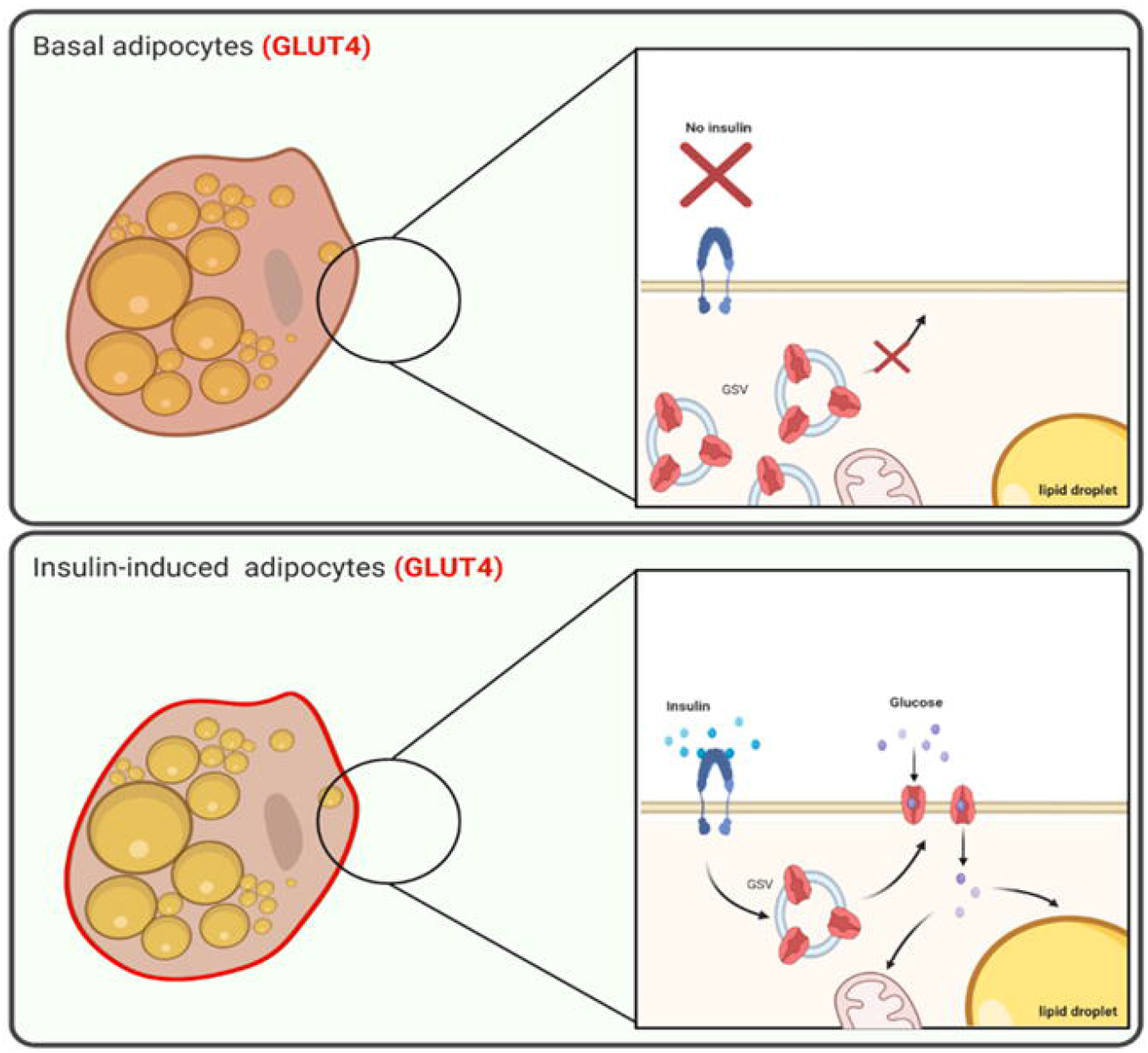
Graphic illustration of the GLUT4 experimental model in adipocytes.

**Fig. 4.**
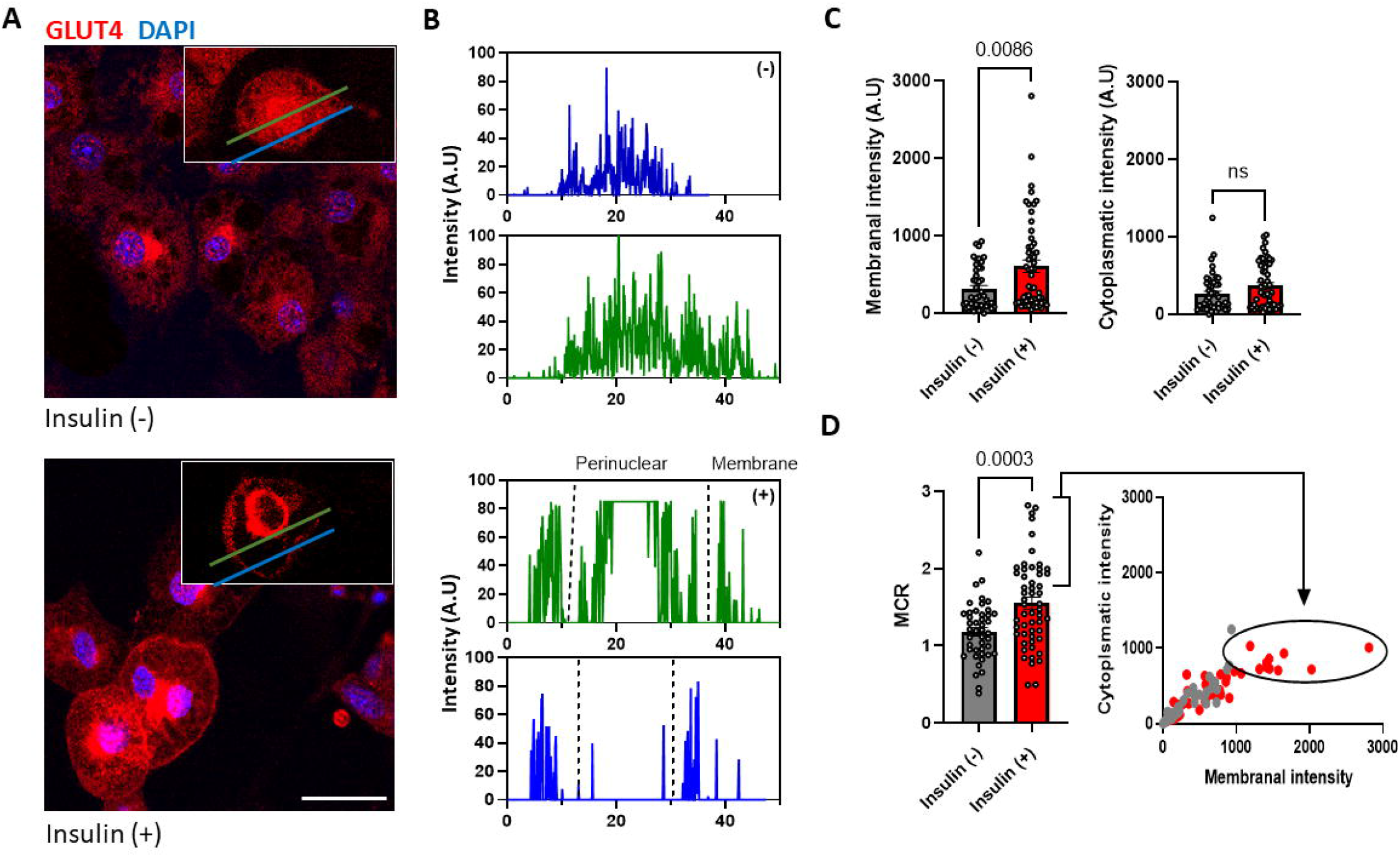
GLUT4 expression and MCR in response to insulin induced in 3T3-L1 cells. (A) Immunofluorescence staining of GLUT4 (red) and Nuclei (blue) in differentiated 3T3-L1 cells (Magnification of X630, scale bar=50μm) (B) Intensity line of the cells’ membrane and cytoplasmic profile. (C) Quantification of the membrane and cytoplasm provide the MCR value of insulin induced (red) and un induced (gray) 3T3-L1 cells, significance was calculated using a two-tailed unpaired student’s t-test, error bars represent means ± SEM.

## Discussion

Here, we present a simple method for quantification of protein translocation between the cytoplasm and PM. The technique is demonstrated by applying the MCR method to monitor the change in GLUT4 expression in insulin induced differentiation of 3T3-L1 cells from pre-adipocytes to adipocytes. The method allows us to compartmentalize the cell and determine the separate levels of protein expression in the cytoplasmic, perinuclear region and PM. Such an analysis can monitor alterations in expression and translocation patterns in response to different cell stimulations. The membrane to cytoplasmatic ratio has previously been used in image-based analysis and biochemical methods^18–21^, but the method described here is novel in that the robust and rapid procedure can be used to quantify the MCR at a single-cell level. In addition, the image-based workflow presented here enables the protein expression and translocation to be quantified in each cellular distribution region.

The quantification of PM-expressed proteins has traditionally been considered a challenge due their complex shape, difficulties in isolation, and overall instability^23,24^. Biochemical assays such as subcellular fractionation followed by immunoblotting or mass spectrometry are often used for quantification although the information obtained is limited due to the batched nature of analysis^24,25^. Imaging of immune stained cells is one of the principal techniques used to quantify membrane expression patterns^26–28^. In comparison to biochemical techniques, IF is appealing because the proteins can be detected directly, in a non-destructive fashion. In addition, the technique is able to provide information at the single-cell level, which is especially important in heterogeneous cellular populations^29^. MCR normalization provides distinct advantages in following the PM and cytoplasm dynamics underlying a cell function, as it is simple to use, reproducible, and comparable. Notwithstanding, MCR analysis has a few limitations as it is used with imaged membranal proteins and the results have to be manually processed although, the accuracy can be improved by using a known membrane protein as both a marker and a normalizer.

Here we demonstrate the efficacy of the MCR method by monitoring the expression and subcellular translocation of GLUT4 in an *in vitro* model of insulin induced adipogenic differentiation of 3T3-L1 cells. While GLUT4 expression in non-induced adipocytes is primarily localized in the perinuclear region, when induced with insulin, the protein is translocated from the perinuclear region to the PM (Fig. 2-4). Accordingly, the MCR analysis indicated an upregulation in membrane expression of GLUT4 after insulin induction. Proper execution of this process is essential for AD function. Our finding that GLUT4 in AD responds to insulin signaling is in agreement with a number of previous reports^12,15,31,32^. Interestingly, some adipocytes were more responsive than others, indicating that molecular activities within a population of cells may be heterogeneous and dynamic, and emphasizing the importance of single-cell-based quantitative methods such as MCR^30,31^. We appreciate that the MCR model may require modifications for other cell types, since in order to assess the parameter values and subcellular distribution patterns, individual morphological extractable attributes must be integrated. However, we predict that MCR quantification will be readily applied to other proteins, and conditions and will be able to reveal hitherto unknown shuttling between cellular compartments.

## Conclusion

In conclusion, the present study describes a simple and accurate approach for protein quantification in the subcellular compartments through immunostaining and MCR calculation. The technique was used on differentiated 3T3-L1 cells induced with insulin to measure the translocation of GLUT4. The results obtained from the presented method demonstrated the upregulation of membrane GLUT4 in response to insulin induction. MCR analysis may pave the way for a better understanding of cellular processes by providing a quantitative method demonstrate the cellular distribution of proteins and enabling us to monitor factors affecting cell metabolism and cellular and molecular signaling. Extensive use of this method in a wide range of conditions will be useful future development of targeted cell therapy and cellular mapping of proteins.

## Acknowledgments

Mike Egozi participated in the project as part of the jInternship program. We acknowledge Ann Avron for the editorial assistance.

